# “Multiphoton holographic photostimulation induces potassium-dependent spike silencing in label-free mouse cortex *in vivo*”

**DOI:** 10.1101/2022.08.06.503049

**Authors:** Stylianos Papaioannou, Johan Zakrisson, Tatiana Kuznetsova, Paolo Medini

**Author notes:** Equal contribution (Stylianos Papaioannou, Johan Zakrisson). Corresponding authorship, <u>Contact:</u> and.

## Abstract

Multiphoton microscopy allows measurement of network activity as well as the manipulation of cell type specific or functionally identified neuronal subpopulations with optogenetic holographic stimulation. When neurons co-express an activity reporter (e.g. calcium or voltage-sensitive indicators) and an (excitatory or inhibitory) opsin, such “all optical” interrogation approaches in vivo allows to draw causal links between function of cell-type specific microcircuits and behaviour. However, the net effects of near-infrared stimulation on network activity per se remain to be adequately investigated in vivo. Here we show that multicell holographic photostimulation with near-infrared radiation with total powers to sample used in current literature halves the spike rate of the non-illuminated neurons in label-free mouse cortex in vivo. The effect is not mediated by GABA release, but depends on NIR-dependent gating of potassium channels as it is absent when neurons are intracellularly perfused with the broad potassium channel blocker cesium ions, and are possibly mediated by heating. The phenomenon may contribute to set an upper limit to holographic photostimulation efficacy, calls for the need to control the effects of holographic stimulation protocols per se in label free preparations, and might be of relevance to interpret the therapeutical effects on infrared stimulation in psychiatry and neurology.

**Highlights:**

- Holographic multi-cell, infrared illumination halves spike rates of no-target neurons
- The effect happens with total powers to sample used in “all-optical” literature
- Infrared-driven spike killing is GABA-independent but depends on potassium channels

## Main text

Optogenetics allows manipulating neuronal activity *in vivo* with high spatio-temporal resolution to draw causal conclusions between microcircuit function and behavior. If neurons co-express an optical sensor (calcium indicator) and an actuator (opsin), multiphoton microscopy allows to simultaneously measure network activity, and photo-stimulate functionally-identified neurons by 3D-holographic optogenetics (by altering light wavefront to split the gaussian beam in several beamlets targeting multiple neurons). Such “all-optical interrogation” of brain microcircuits is the frontier in system neuroscience (e.g.[1, 2]). Recent approaches increased phostimulated neurons to hundreds[3–5], implying brain illumination with absolute power to whole sample: 0,4-4 W). However, the net effect of near-infrared-radiation (NIR) on brain network *in vivo* remains largely unexplored and controversial, as some reports show inhibitory (e.g.[6]), while other excitatory (e.g.[7]) effects. This knowledge is needed to optimize/interpret all-optical interrogation studies, but also promising NIR-based therapies in psychiatry (e.g.[8]).

We addressed this by using 3D holographic stimulation with NIR using conditions previously implemented by others (1040 nm, 10 neurons photostimulated scanless for 500 ms, 16x objective, 45 mW/neuron) while performing two-photon-assisted(tpa)-*in vivo* whole-cell recordings with “shadowpatching” technique from one not-illuminated neuron in the photostimulated network in mouse parietal cortex (see Supplementary Materials). As the example and peristimulus-time-histograms (PSTH) spike counts show (Figure 1A, left and middle plots, respectively), we observed a clear, ca 2-fold reduction of firing rates locked with NIR illumination. Spike rate drops were quantified as their Z-score modulations, to normalize and compare the activity of cells with different spontaneous activity levels and variability of temporal spike patterns (see Methods). Figure 1A, rightmost plot, shows the grand-averaged Z-score with 5-95 confidence interval and its clear negative modulation during NIR-illumination (means before and during illumination: −0.121±0.016 vs −0.989±0.591; N=6 mice, n=10 neurons; t=2.477, p_2-tail_=0.038).

**Figure 1.**
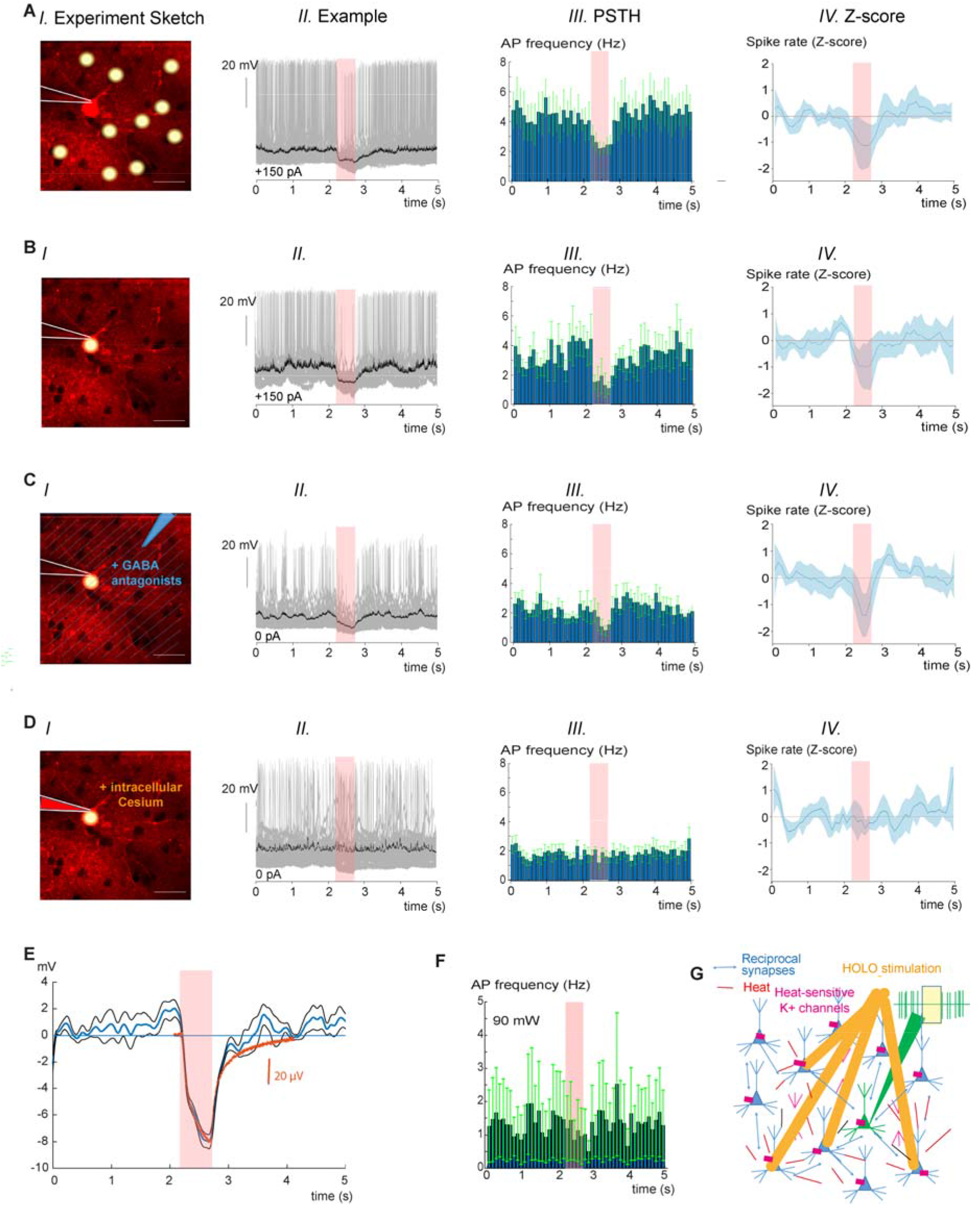
NIR-holographic illumination evokes a non-GABAergic, potassium-dependent spike reduction during both on-target and multi-off target stimulation in mouse cortex *in vivo*. **A**. *Multi-target holographic NIR illumination causes a laser-locked spike reduction in non-illuminated neurons*. From the left to right, respectively: ***I*.** <u>Experimental sketch</u> showing the *in vivo*, two photon assisted shadow patch of one neuron while 10 target cells were holographically illuminated in the intervention volume – calibration bar: 50 μm; ***II*.** <u>Example</u> showing 50 overlaid traces (pink rectangle represents the illumination time, 500 ms). 150 pA of background current was injected. In all examples the overlaid single traces are in gray, whereas the averaged trace is in black; ***III*.** Population <u>PSTH</u> of spike counts for this experimental groups (n=10, where n is number of cells; mean firing frequencies without and with illumination, respectively: 4.661 ± 4.785 vs 2.503 ± 0.792; t-test paired: t = −2.217, p_1-tail_ = 0.026); ***IV*.** Grand average Z score of spike counts (blue line; light blue area is the 5-95 confidence intervals, see statistics in the main text). **B**. *On-target, full power holographic NIR illumination causes a laser-locked spike reduction*. ***I*.** Experiment sketch. ***II*.** Example showing 20 overlaid traces (red line represents the illumination time, 500 ms). A background current injection of 150 pA was done. ***III*.** Population PSTH of spike counts for this experimental groups (n=7; mean firing frequency without and with illumination, respectively: 3.613 ± 6.070 vs 1.494 ± 2.358 Hz; t-test paired: t =-2.625, p_1-tail_ = 0,019). ***IV*.** Grand average <u>Z score</u> of spike counts (blue line; light blue area is the 5-95 confidence intervals, statistics in main text). **C**. *On-target, full power holographic NIR illumination causes a laser-locked spike reduction also upon extracellular GABA-A and GABA-B receptor blockade*. ***I*.** Experiment sketch). ***II*.** <u>Example</u> showing 30 overlaid traces (pink rectangle represents the illumination time, 500 ms). ***III*.** Population <u>PSTH</u> of spike counts for this experimental groups (n = 7 neurons; mean firing frequency without and with illumination, respectively: 2.258 ± 1.539 vs 1.578 ± 1.789; t-test paired: t = −4.251, p_1-tail_ = 0.001). ***IV*.** Grand average <u>Z score</u> of spike counts (blue line; light blue area is the 5-95 confidence intervals, statistics in main text). **D**. *Potassium channels blockade by intracellular perfusion with cesium of the holographically illuminated neuron prevents the NIR-driven spike reduction*. ***I*.** Experiment sketch. ***II***. <u>Example</u> showing 23 overlaid traces (pink rectangle represents the illumination time, 500 ms. ***III*.** Population <u>PSTH</u> of spike counts for this experimental groups (n = 5 neurons; mean firing frequency without and with illumination, respectively: 1.886 ± 0.751 vs 1.666 ± 0.362; t-test paired: t = −1,173, p_1-tail_ = 0.153). ***IV***. Average <u>Z score</u> of spike counts (blue line; light blue area is the 5-95 confidence intervals, statistics in main text). **E.** *Subthreshold effect of NIR illumination indicates a decrease of membrane resistance*. Grand averaged subthreshold (membrane potential) response to full power irradiation of intracellularly recorded neurons. A positive current of 150 pA was injected as background, implying that the relative hyperpolarization observed reflects a NIR-induced membrane resistance drop. The blue curve is the grand-average and the black curves show 5-95 confidence intervals. Note the similarity of the time course of normalized resistance drops of a pipette in the bath (orange trace, normalized to the minimum peak) with that of neurons *in vivo*. Traces are corrected for the pipette and access resistance drops, respectively for *in vitro* and *in vivo*, due to irradiation. **F.** *Lack of spike silencing when a single neuron is irradiated with 90 mW* (no background current injection: 1.325 ± 2.570 vs 1.112 ± 1.807, t-test paired, t=-1.202, p_1-tail_ = 0.148). **G.** *Model representing that NIR reduce spiking via potassium-channel gating*. The results of E and F, the uniformity of the response to NIR across neurons, its quick temporal kinetics, together with the similarity of the previous subthreshold plots with the temporal dynamics of temperature increase of the brain parenchyma in our experimental conditions (as of[13]), raise the hypothesis that NIR during multi-target, holographic photostimulation acts via heat gating shunting potassium channels, with a negligible effect via synaptic network.

To investigate the mechanism of NIR-induced spike killing, we performed single-cell holographic stimulation with full power on single neurons (450 mW). Although this power level is ca 4 times higher compared to traditional protocols *in vivo* [1, 2] with spiral-scanned illumination of somatas, the power density (mW/μm^2^) had similar value to our scanless approach. The reduction of spiking was similar as seen in the example and population PSTH, and in the Z-score grand-average (Fig. 1B, mean Z-scores before and during illumination: - 0.244±0.055 vs −1.077±0.301; N=5 mice, n=7 neurons; t=2.661, p_2-tail_=0.037).

We tested first whether the NIR-induced spike silencing could be due to photo-induced GABA release [9]. We repeated tpa-patch and holography experiments in presence of subepileptic dosages of GABAA and GABAB blockers (see Methods). The example, population PSTH and Z-score plot confirm that the NIR holographic light pulse induces a clear spike reduction also when GABAergic transmission is blocked (Fig. 1C; mean Z scores before and during illumination: −0.185±0,037 vs −1.146±0,453; N=4 mice, n=7 neurons, t=2.988, p_2-tail_=0.026).

Having excluded the role of GABA-_A_ (chloride-dependent) and GABA-_B_ (potassiumdependent) mechanisms, the possibility remains that NIR light directly activates potassium conductances, capable of hyperpolarizing and shunting the neuron. Such a mechanism is suggested by observing NIR-induces membrane voltage drop of 7.0±0.4 mV upon +150pA sustained current injection, what should correspond to ca 50% reduction of membrane resistance (Figure 1E, left). We thus performed tpa-patch and holography filling pipettes with Cs^+^ ions to dialyze intracellular potassium and block many potassium channels. After physiologically effective intracellular perfusion with Cs^+^ (half-width-at-half-height of spikes>5 ms, see Methods), we performed holographic stimulation on the patched neuron. As the example, population PSTHs, and the grand average Z score plot show (Figure 1D), holographic illumination *failed* to induce spike silencing in cesium-perfused neurons (mean Z scores before and during illumination: −0.023±0.038 vs −0.184±0.104; N=5 mice, n=5 neurons, t=1.003, p_2-tail_=0,372).

Thus, NIR, network holographic photostimulation of label-free cortex in vivo halves the spike output of not-illuminated neurons largely via gating of potassium-selective channels. Similar or higher power/cell (up to 90-100 mW) have been used in holographic manipulations[1, 2, 10]. New lasers with higher peak powers and slower repetition rates allow stimulation with 10-fold less power/neuron. However when ≥10-fold more neurons are stimulated (as e.g. [4, 5]), the total power, delivered to holographic intervention volume, is comparable [5] or higher [10],[4] than ours. NIR-induced spike reductions *might contribute to set an upper limit to photostimulation efficacy* in terms of max_spikes/neuron[1] or percentage of activated cells, which is *smaller* for higher powers[5]. The latter observation, and activity reductions related to total power during photoexcitation ([11]-Figure 3), have been attributed to GABAergic network activation – which we excluded. Our data suggests instead a direct role of NIR-induced potassium-channels gating in these observed inhibitory effects in non-targeted neurons. The fact that we observe photoinhibition upon multi-cellular holographic stimulation in non-illuminated cells (to a similar extent as in on-target experiments) suggests the contribution of non-light, heat-dependent effects on the observed phenomenon (but see[12]). The lack of effect when a single neuron was targeted at 90 mW (Figure 1F) excludes the role of withdrawal of excitation from neighbouring pyramids. The same is also proven by the *net* membrane conductance increase, instead of decrease. NIR-locked potassium channels gating is possibly mediated by *heat* as suggested by (1) the similarity between the temporal dynamic of voltage (resistance) drop of neurons *in vivo* and the solely illuminated pipette, and (2) the simulated temperature increase in our conditions ([13]; see Figure 1E, orange trace; though recent report of midIR-driven, selective photochemical gating of potassium channels[14]).

Our results are possibly behaviourally-relevant because animals detect sparse activity modulations[11]. We used continuous illumination (500 ms), whereas others use shorter illumination to drive faster opsins, which can affect heat delivery[13]. However, to drive/modulate behavior, total illumination times have to last hundreds of ms or seconds (e.g.[3]). Ultimately, although the impact of our findings might *vary quantitatively* in different holographic protocols (using opsins with different kinetics (e.g.[1, 5] vs[3, 4, 10]), our results prompt to design holographic photostimulation so to *minimize total power* and/or *control for such effects* for data interpretation (see[2],Figure 5). Further studies will characterize whether and how more prolonged NIR illumination modulate network activity dynamics, such as during two-photon network imaging or NIR-based photostimulation therapies effective in psychiatry[8].

## Abbreviations

NIR: near-infrared radiation
PSTH: peristimulus time histograms
tpa: two-photon assisted

## Author contributions

S.P and P.M. conceived the study and designed the experiments. S.P, J.Z. and T.K gave similar contribution to experiments, J.Z. characterized and maintained in service the holographic system in discussion with and with inputs from S.P and P.M.. S.P. designed the electrophysiology analysis routines. Data analysis was done by S.P. and J.Z, and T.K. contributed to dataset organization and example selections. P.M. coordinated and supervised all the work and wrote the manuscript. All author contributed with critical inputs to the manuscript and approved its final version.

## Acknowledgments

We thank Per Utsi for excellent technical assistance, Sebastian Sulis Sato for help in molecular construct design and participation in initial experiments, József Orbán from Femtonics for assistance on the use of the holographic module and Marco Dal Maschio for technical inputs and scientific discussions.

## Declaration of competing interests

The Authors declare no competing interests

## Supplementary Materials and Methods

### 1. Holographic 2-photon stimulation

The microscope (Femto2D, Femtonics, Budapest, Hungary) used for all two photon (2P) experiments in this paper was composed of two main parts: a 3D holographic module and a 2P imaging module. The holography module used for photostimulation, designed by and purchased from Femtonics, was driven by the following source: Femto train 1040-5, fixed wavelength: 1040 nm, pulse duration: 220 fs, pulse frequency: 10 Mhz, Spectra-Physics. A reflective spatial light modulator (SLM, Hamamatsu: X13138-03 LCOS-SLM 1272×1024 pix, 12.5 μm size, 15.9×12.8 mm chip size, refresh rate 60 Hz) was used to generate the stimulation pattern. Femtonics provided a computer running a dedicated software to generate and then display the phase mask on the SLM via a dedicated graphic card connected to the SLM driver electronics via DVI connector with a refresh rate of 60 Hz. The control software provided a loo the zero order component depending on mask complexity -). The phase masks to be displayed on the SLM were calculated with a weighted Gerchberg-Saxton algorithm (Gerchberg, R. W.; Saxton, W. O., 1972. “A practical algorithm for the determination of the phase from image and diffraction plane pictures”. Optik. 35: 237–246). The weighting partially compensated for the diffraction efficiency at targets at the periphery of field of view was lower compared to targets in the center (an effect possibly due to increasing interpixel crosstalk at larger diffraction angles [1]). The holographic intervention field of view had 50% linear size compared to the imaging field of view through the objective. A schematic of the microscope as provided by Femtonics is drawn in Supplementary Figure 1A. The source of the 2P fluorescence imaging was a tunable pulsed NIR laser (Chameleon Discovery, wavelength range: 700-1300 nm, pulse width: 160 fs, pulse frequency: 80 Mhz, Coherent). The two beams, i.e., the imaging laser and the photostimulation (holography) laser, are combined using a dichroic mirror (“Lightpath mirror” in Supplementary Figure 1), and sent through a 16 ×(0.8 NA) water immersion objective (Olympus). The registration and alignment of the photostimulation and imaging paths was achieved by using a widefield camera that imaged the sample via objective and tube lens while phostostimulating a homogeneously fluorescent sample (as exemplified in[2]).

Experiments were done in brain areas without major blood vessel and adequate transparency until a depth of 500 μm. The holographic photostimulation was controlled via an electro-mechanical shutter driven by the electrophysiology software (Patchmaster, HEKA, Germany). Neurons were stimulated using a 15 μm diameter circle (corresponding to a power density of 2,54 mW/μm^2^ when full power was used on a cell, or 0,25 mW/ μm^2^ – corresponding to 45 mW/neuron - when split onto 10 targets). For initial calibration of the system, we measured the point spread function (psf) of the holographic system illuminating at full power a 15 μm circular disk in a fluorescein solution by imaging the fluorescence intensity via the imaging path (see Supplementary Figure 1B). To measure whether our holographic system has the capacity to achieve holographic stimulation *in vivo* with single cell resolution, we infected mice pyramidal cells with adeno-associate viruses coding for the excitatory opsin bReaches, and the YFP fluorescent markers. So, we could perform two-photon targeted patch clamp (relaying on fluorescent marker expression) and holographic illumination of single neurons expressing the opsin bReaches[3]. In that case, green fluorescent pipettes filled with Alexa 488 were used to perform in vivo two-photon targeted patch clamp ([4] and see Supplementary Figure 1C, left). For these excitatory holographic experiments the power per cell was 90 mW (similar to the one used by[1] or[5], although in their case a diffraction limited beam was spiral-scanned on the neuronal somata, in contrast to our scanless approach which was similar to the one used *in vivo* by e.g.[6]).

We next documented that our holographic system was able to perform multiphoton holographic excitation with single cell resolution *in vivo*. As shown in Supplementary Figure 1C, right traces, *in vivo* two-photon targeted patch of bReaches expressing cells shows that suprathreshold excitation (reliable action potential firing in response to every stimulation) was obtained when the beam was centered onto the soma of the recorded neuron, but not when it was displaced 30-40 μm laterally (circa two somata diameter; N=2 mice, n=3 neurons).

### 2. Two-photon targeted in vivo whole-cell recordings

Whole-cell current-clamp recordings were acquired using EPC 10 USB Quadro Patch Clamp Amplifier integrated with LIH 8+8 data acquisition interface and the Patchmaster software (HEKA, Germany). Signals were 1000x amplified and digitized at 50 kHz. Data acquisition was triggered by software that was controlling the holographic photostimulation (MES; Femtonics, Hungary). Glass micropipette (6-8 MΩ; Ø 1 μm) filled with intracellular solution containing Alexa 594 dye (Sigma, 20 μM) for visualization was positioned with a 45 deg angle under the coverslip glass in L2/3 of somatosensory cortex for subsequent 2-photon targeted patch-clamp following the “shadowpatching” method in label-free cortex[7]. Unless otherwise stated, the standard intracellular solution was composed as follows (in mM): 135 K-gluconate, 10 Na2-phosphocreatine, 4 KCl, 4 ATP-Mg, 0.3 GTP-Na (hydrate), 10 HEPES; adjusted with KOH to pH=7.2).

In a series of experiments Cs^+^-based intracellular solution was used in order to block potassium channels, where K-gluconate and KCl from standard intracellular solution was substituted by Cs-methanesulfonate, and CsCl, respectively. In the experiments with Cs^+^- based intracellular solution we considered the potassium channels to be successfully blocked when the spike width was becoming broader than 5 ms (the width was calculated as the fullwidth at half-maximum value of the action potential peak relative to the pre-spike baseline membrane potential), which happened minimum after 10-15 minutes from achieving a wholecell configuration.

In order to block GABAergic transmission the GABAA and GABAB receptors’ antagonists gabazine (SR 95531, Sigma-Aldrich) and CGP52432 (Sigma-Aldrich) were applied in the bath at sub-epileptic concentrations (1.5 μM and 1 μM, respectively; see [8]) at least 15 min before start of acquisition. The effectiveness of drugs was proven by the modifications of spontaneous activity as observed in [8].

Recordings started when the somata of patched cell was getting filled with the Alexa 488 dye to ensure its position so to guide positioning the hologram. In experimental series without pharmacological manipulations (extracellular GABA blockade or intracellular Cs^+^ perfusion), a background current of 150 pA was injected to drive neurons to spike, due to the spike sparsness in layer 2/3 pyramidal neurons (e.g.[9]). Each series of recordings contained 20-100 sweeps (depending on the level of spiking activity), and each sweep lasted 5 s. During 2.2-2.7 s of each sweep holographic photostimulation (500 ms illumination time, 1040 nm excitation wavelenght, 450 – or 90 mW average power) was delivered. The stimulation was either single-spot (Ø 15 μm) placed on the patched cell, or multi-spot with 10 targets (Ø 12-15 μm each) distributed in 300×300 μm plane, each placed at least 30 μm away from the patched cell. Input resistance was monitored during the experiments by passing series of square-wave (-or+100pA) current pulses to hyperpolarize or depolarize the patched cell and varied between 25 and 75 MΩ (variation larger than 100% were a criterion for rejection). Only recordings with stably polarized membrane potential and clearly visible spikes were included.

### 3. Animal procedures

#### - Animals

Young (6 - 11 weeks) wild-type C57BL/6J male mice were used. They were housed at Umeå Center for Comparative Biology’s (UCCB) and maintained in a 12 h light/dark cycle, with food and water available ad libitum. At the end of the experiment animals were euthanized by isoflurane overdose followed by cervical dislocation. All procedures were done according to the Swedish Ethical Committee for Northern Sweden (permission number A26-2018).

#### - Viral injections

To validate the efficacy of our holographic system, we expressed excitatory opsin bReaches in pyramidal neurons located in L2/3 somatosensory cortex of mice. For that 6-7 weeks old wild-type C57BL/6J mice were injected with 750 nl viral mix (AAV serotype 8, produced by AAV Core Facility, Lund, Sweden), containing bReaches-YFP-floxed + CaMKII-CRE in proportion 1:0.5 (titre: 10e13/mL). A red fluorescent, Alexa 594-filled patch pipette was used to perform dual-color two-photon in vivo targeted patch – see Supplementary Figure 1C, left). To achieve expression in somatosensory cortex and avoid the scar tissue above the expression site, the injection pipette was inserted at 45° to horizontal plane and through a tiny craniotomy (-1 mm on antero-posterior, and 2 mm on medio-lateral from bregma) at a depth of 700 μm. Recordings were done from 6 weeks to 3 months after the injections.

#### - Surgery

For in vivo 2-photon targeted patch-clamp recordings combined with holographic stimulation, wild-type mice were implanted with a headbar (for head-fixation) and operated to insert half-circular optical window (for imaging and holographic stimulation) with access for glass micropipette (for patch-clamp) to the parietal cortex. Animals were anesthetized with isoflurane (4,5% for induction; 1.5-2.5% during surgery). To reduce the inflammatory response Dexamethasone (0.67 mg/kg, i.m.; 60-100 μl of 0.2 mg/ml in saline) was injected i.m. before opening the cranial window. Body temperature during the experiments was constantly monitored with a rectal thermocouple probe and maintained at 37 °C with a heating plate (using ATC-2000 Animal Temperature Controller). After removing the skin and periosteum, the metal head-plate was glued to the bone with Superglue and tissue glue (3M Vetbond) in the hemisphere opposite to the planned craniotomy. Using the head-plate animal was mounted on a custom-made stereotaxic holder. Then a bath chamber was built with dental cement (Paladur) and formed at the borders of the skull and reinforced by tissue glue. The bath allowed to keep the cortical surface always covered by physiological solution (0.9% NaCl) and to place the ground electrode. After the bath chamber, the parietal bone plate was carefully thinned by tangential drilling, and big rectangular craniotomy was cut above somatosensory cortex. Then half of the craniotomy was covered with the half of the 5 mm cover glass (to allow the imaging and holographic stimulation), leaving another half of the craniotomy exposed for pipet access. The glass was glued to the bone with Superglue.

### 4. Data analysis

#### - Suprathreshold data

Action potential rates were analyzed in MATLAB R2019b. An action potential was defined as a positive deflection crossing the threshold of > 5 SD of the mean first derivative in time of the Vm. The sweeps were included into the analysis if the resting membrane potential was ≤-30 mV (including Cs+ experiments), without correcting for liquid junction potentials and offset through access resistance. Z-scores were computed by subtracting the mean pre-illumination (spontaneous) activity in each trial and by dividing each value with the standard deviation of the spontaneous activity, and the z-score sweeps were averaged for each recorded neuron. The grand average of Z scores are presented in all the rightmost panels of Fig 1 for each experimental groups (thick blue lines), and 5-95% confidence intervals are shown by the light blue shaded areas. For statistical comparison of Z-score modulation, the Z values of each cell of a pre-stimulus and of a steady-state intrastimulus epochs of same duration were defined (300 msec) and compared with a paired statistics. PSTHs of spike activity are shown with 100 ms binning. For statistical comparison of the absolute firing rates, the spontaneous firing rate (in absence of stimulation) was compared to the mean firing rate during IR illumination via paired t-tests.

#### - Subthreshold data

For the analysis of the subthreshold response to NIR holographic illumination, action potential were removed after detection described as above and the Vm was replaced by linear interpolation, and the sweeps were averaged. The baseline was set to zero and the hyperpolarization amplitude was measured. According to our observation which is in line with previous published work (e.g. https://knowledge.reagecon.com/wp-content/uploads/2021/09/Technical-Paper_The-Effect-of-Temperature-on-Conductivity-Measurement.pdf), NIR illumination on the glass pipette/silver electrode complex, creates a drop of the pipette resistance due to photochemical as well as heating effects. This resistance and subsequent voltage drop needs to be calculated and subtracted from the whole cell in vivo recordings so as the residual voltage drop can be attributed to a net NIR effect on the cells and not to any change on the pipette or access resistance. To implement that first we did measurements (in current clamp configuration) of the voltage drop of the pipette only under the same intensity of NIR illumination as we kept during the experiments (450 mW). Based on these measurements we calculated the voltage drop based not only on the pipette but also on the access resistance as measured after the whole cell access. On average the above effect accounted for a 6.2% voltage drop during whole cell *in vivo* measurements.

## Supplementary Figure and Legend

**Supplementary Figure 1.**
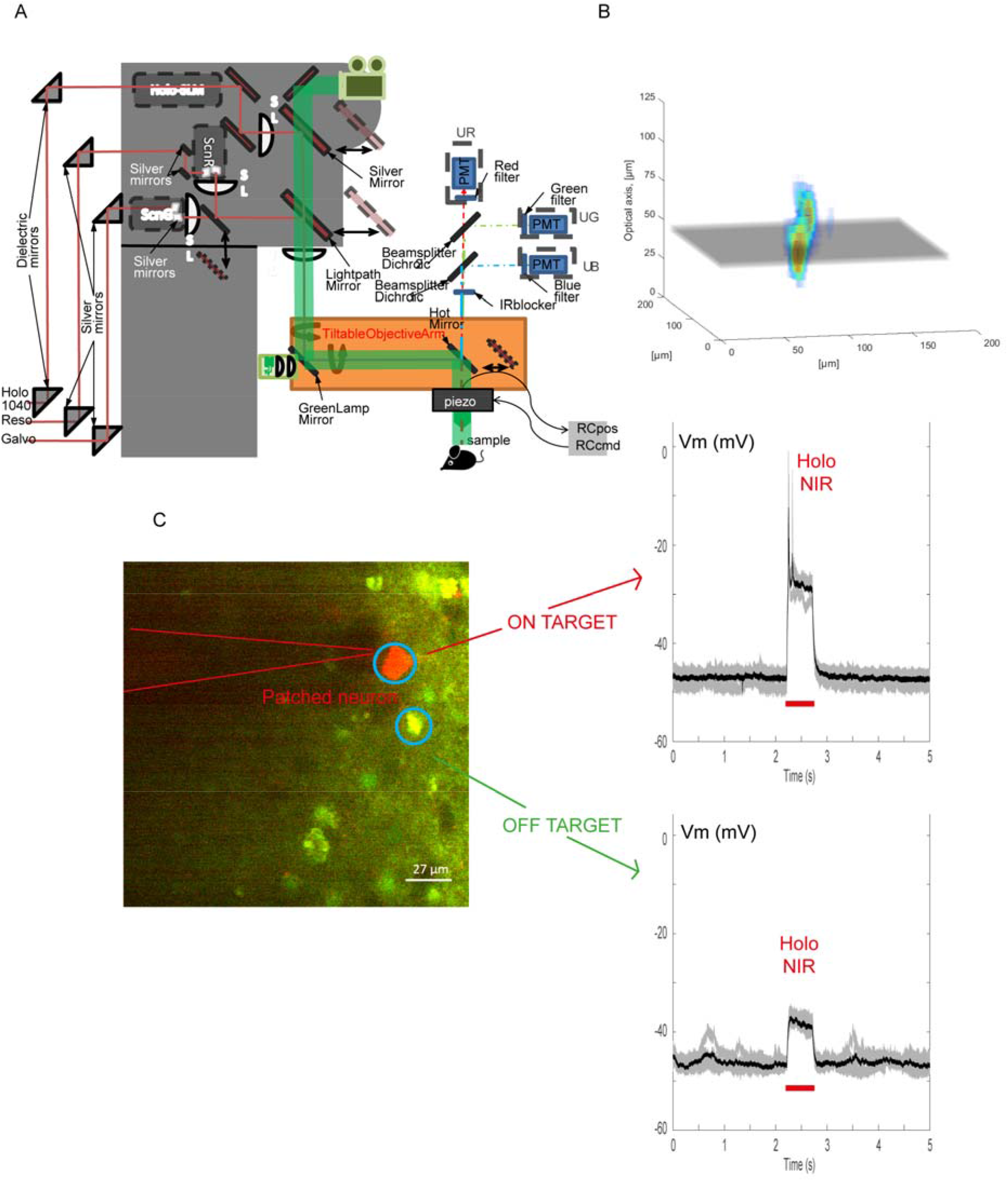
Holographic set up characterization and photostimulation *in vivo*. **A**. *Inner schematic of the two-photon microscope* as kindly provided by Femtonics. ScnG: galvanometric scan mirrors; ScnR: resonant (30 Hz) scan mirrors; SL: scan lens; SLM: spatial light modulator; PMT (UG, UB, UR): photomultipliers for green, blue and red, respectively. **B**. *Visualization of the holographic system’s optical psf* as obtained by imaging the excitation volume of a uniform fluorescein filled obtained by projecting the full power hologram (15 μm circle), the same the one as used in the on-target, full-power experiments (as in Fig 1 B, C, D). The gray plane is the focal plane. **C, D.** *Validation of the physiological psf: demonstration of holographic photoactivation of bReaches-expressing cortical neurons in vivo with single cell resolution*. **C** shows a bi-color image of cortical neurons expressing the green fluorescent protein YFP in cytoplasm and bReaches in the membrane, while the pipette tip was visualized under two-photon microscopy (pipette was filled with the red fluorescent indicator Alexa 594 (20 μM)). **D** shows overlaid single sweeps (voltage traces in current clamp) upon phostimulation (90 mW, 500 ms, 1040 nm) of the patched neuron (top trace, red arrow) or of a neighboring neuron with soma located within 30 μm from the recorded neuron (bottom trace, green arrow). In these examples the overlaid traces are in gray, whereas the averaged traces are in black.

## Notes

### Competing Interest Statement

The authors have declared no competing interest.

